# Hidden from plain sight: Novel *Chlamydiota* diversity emerging from screening genomic and metagenomic data

**DOI:** 10.1101/2023.03.17.533158

**Authors:** Helen R. Davison, Gregory D.D. Hurst

## Abstract

*Chlamydiota* are an ancient and hyperdiverse Phylum of obligate intracellular bacteria. The best characterized representatives are pathogens or parasites of mammals, but it is thought that their most common hosts are microeukaryotes like Amoebozoa. The diversity in taxonomy, evolution, and function of non-pathogenic *Chlamydiota* are slowly being described. Here we use data mining techniques and genomic analysis to extend our current knowledge of *Chlamydiota* diversity and its hosts, in particular the Order *Parachlamydiales*. We extract one *Rhabdochlamydiaceae* and three *Simkaniaceae* genomes from NCBI Short Read Archive deposits of ciliate and algal genome sequencing projects. We then use these to identify a further 14 and 8 genomes respectively amongst existing, unidentified environmental assemblies. From these data we identify two novel clades with host associated data, for which we propose the names ‘*Candidatus* Sacchlamydia’ (Family *Rhabdochlamydiaceae)* and ‘*Candidatus* Amphrikania’ (Family *Simkaniaceae*), as well as a third new clade of environmental MAGs ‘*Candidatus* Acheromydia’ (Family *Rhabdochlamydiaceae*). The extent of uncharacterized diversity within the *Rhabdochlamydiaceae* and *Simkaniaceae* is indicated by 16 of the 22 MAGs being evolutionarily distant from currently characterised genera. Within our limited data, we observe great predicted diversity in *Parachlamydiales* metabolism and evolution, including the potential for metabolic and defensive symbioses as well as pathogenicity. These data provide an imperative to link genomic diversity in metagenomics data to their associated eukaryotic host, and to develop onward understanding of the functional significance of symbiosis with this hyperdiverse clade.

**Graphical Abstract:** 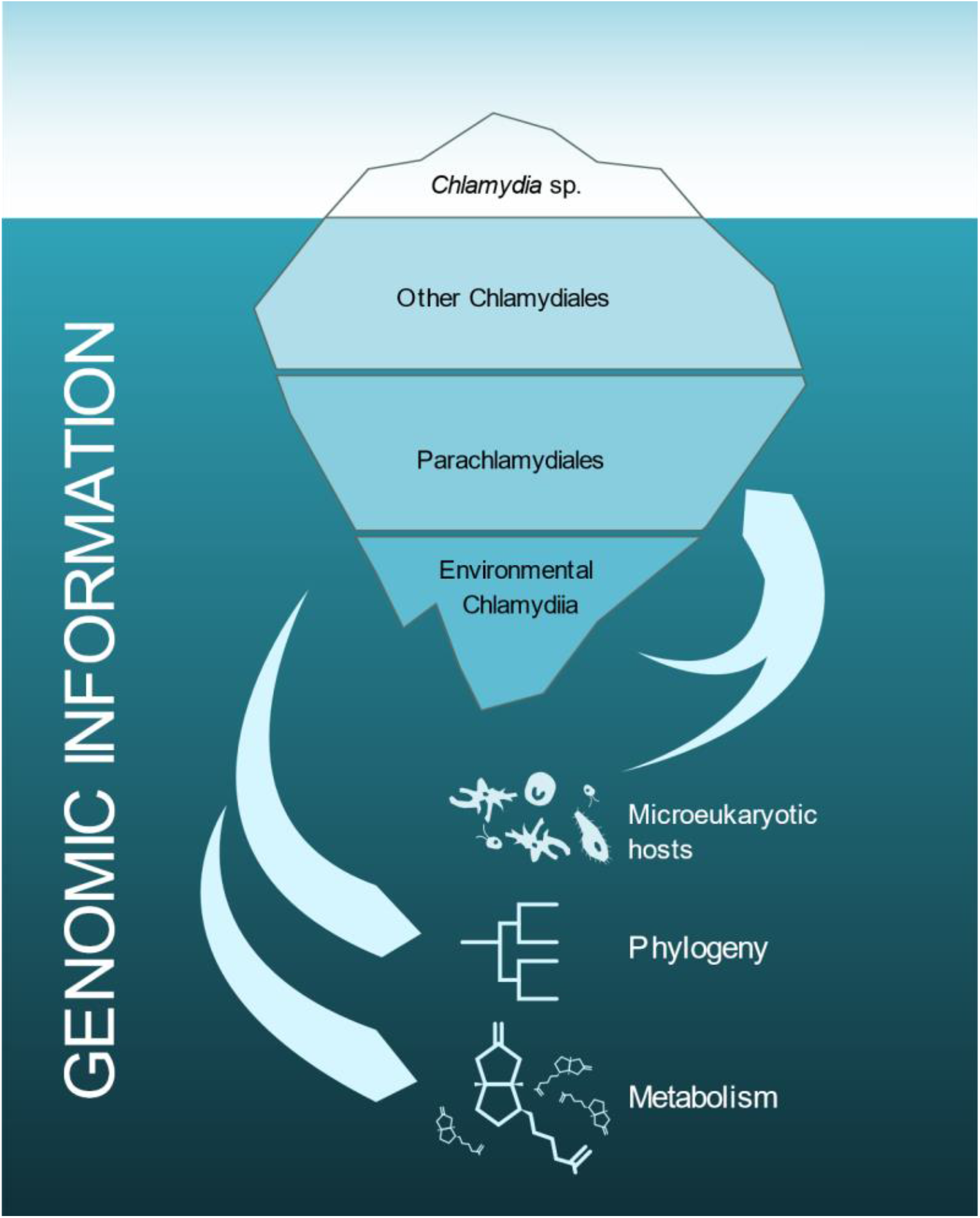

## Introduction

*Chlamydiota* are a phylum of bacteria that are highly diverged from any other microbial clade, such that they were for some time considered a separate kingdom of life (Prowazek, 1912; Horn, 2008). Species within the group uniquely share a biphasic lifestyle, with an inert non-dividing environmental stage and a reproductive intracellular stage (Kahane *et al*., 1993; Bastidas *et al*., 2013; König *et al*., 2017). These bacteria also share proteins that are associated with chloroplast function in plants, which has been speculated to suggest a long-standing relationship with cyanobacteria and algae (Brinkman *et al*., 2002; Horn *et al*., 2004). It has also been proposed that ancient members of *Chlamydiota* facilitated the establishment of the first photosynthetic proto-algal eukaryotes (Huang and Gogarten, 2007).

*Chlamydiota* are most well known as mammalian pathogens and were first described by the ancient Chinese and Egyptians (Gruber, Lipozenčić and Kehler, 2015). However, the status of the clade has been revised since the advent of DNA taxonomy and it is now recognised that their host range is much broader. They are commonly found in free living amoeba species and environmental samples, though here the original host is often not known (Horn, 2008). Besides amoebae, they can also infect mammals, fish, worms, and a variety of arthropods (Draghi *et al*., 2004; Horn, 2008; Kjeldsen *et al*., 2010; Halter *et al*., 2022). There are potentially many hundreds of unexplored families within *Chlamydiota* (Horn, 2008; Lagkouvardos *et al*., 2014; König *et al*., 2017). One order of *Chlamydiota*, the *Parachlamydiales*, are widely observed and sequenced in environmental samples, but have little to no functional information and recognised taxonomic affiliations are currently fluid (Horn, 2008; Nylund *et al*., 2018; Halter *et al*., 2022).

Very recently, *Parachlamydia acanthamoebae* has been shown to protect its amoeba host from giant viruses (Arthofer *et al*., 2022). The mechanism for this defensive symbiosis is not yet known, but *P. acanthamoebae* seems to prevent the virus from co-opting the amoeba machinery, thus preventing the formation of viral factories. Additionally, *Neochlamydia* has previously been implicated in defensive symbioses against *Legionella pneumophila* (Ishida *et al*., 2014; Maita *et al*., 2018). These data indicate the potential these symbioses have as hidden players in the biology, ecology, and evolution of microeukaryotes beyond a role as pathogens. Moreover, this strong impact on host biology creates an imperative to investigate symbioses in microeukaryotes outside of amoebae.

Sequence data provides an opportunity to estimate the level of uncharacterised diversity in microbial symbionts. Two approaches are possible. First, genome sequencing projects for eukaryotes commonly contain the reads from their associated symbionts within them. Second, environmental sequencing projects allow the identification and assembly of draft genomes without host association data. Here, we first screen the NCBI Sequence Read Archive (SRA) databases for ciliates, macro- and micro-algae for the presence of *Chlamydiota* and assemble 1 and 3 new draft genomes for the families, *Rhabdochlamydiaceae* and *Simkaniaceae* respectively (both *Parachlamydiales*). We then use these genomes to identify 22 related MAGs from environmental metagenomic studies on online databases. Finally, we conducted a phylogenetic and metabolic analyses of these, and existing genomes, to assess their taxonomic placement and explore the potential impacts on their hosts.

## Materials and methods

### Data collection

The presence of *Chlamydiota* bacteria was investigated across Algae and Ciliate SRA databases on NCBI as described previously in Davison et al. (2022) using PhyloFlash to identify deposits with evidence of Chlamydiota bacteria 16S rRNA sequence. Those SRA providing evidence of *Chlamydiota* presence were taken forward to metagenomic binning and assembly. We used GTDBtk to screen bins and found that all genomes recovered were related to *Rhabdochlamydiaceae* or *Simkaniaceae*, so analyses from this point on were focused to Parachlamydiales. Genomes assembled here were then used as baits to identify related metagenomically assembled genomes (MAGs) in the NCBI nr database that were previously not fully classified.

Existing *Parachalmydiales* genomes, as well as a *Clavichalmydia* and a *Chlamydia* genome were downloaded from NCBI (see Supplementary data 1) alongside the GEM catalogue for genomes identified by Köstlbacher et al. (Köstlbacher *et al*., 2021; Nayfach *et al*., 2021).

### Metagenomic assembly and annotation

Minimap2, MEGAHIT and MetaBAT2 (Li *et al*., 2015; Li, 2018; Kang *et al*., 2019) were used to assemble and bin metagenomic bacterial genomes from SRA data using the pipeline in Davison (2022). Genome quality and identities were checked with CheckM and NCBI Blastn (Camacho *et al*., 2009; Parks *et al*., 2015).

Four newly assembled genomes, 18 environmental MAGs, 82 previously established *Parachlamydiaceae*, and 2 outgroup *Chlamydiaceae* were then passed to Anvio-7 to be annotated with COG20 and KEGG kofams through their pangenomics pipeline (Aramaki *et al*., 2020; Eren *et al*., 2021; Galperin *et al*., 2021; Davison, 2022).

### Phlyogenomics and metabolic predictions

Clusters of protein homologs shared by 90% of the 106 genomes used were identified and extracted through Anvio-7. We extracted 34 single copy core gene clusters that contain a total of 3604 genes. A phylogeny partitioned by gene cluster was constructed with IQTREE, using model finder plus, 1000 ultrafast bootstraps, and 1000 SH-alrt replicates. Models per partition can be found in Supplementary data 2.

We used GTDBtk for taxonomic classification of the recovered genomes and to identify taxonomic novelty among them. AAI was calculated for all genomes through kostalabs online enveomics suite (Rodriguez-R and Konstantinidis, 2016). ANIb was calculated with pyANI through anvio-7 (Supplementary data 1) (Pritchard *et al*., 2016). The network was visualised in Gephi 0.9 (Bastian, Heymann and Jacomy, 2009), with edges filtered for >0.6 AAI and >0.95 ANI scores. Annotations were added in Inkscape (Inkscape Project, 2020).

Anvio-7 using KEGG kofams was used to assess metabolic completeness for high quality genomes exceeding a CheckM score of 90% completeness to minimize the loss of information caused by incompleteness (Eisenhofer, Odriozola and Alberdi, 2023). Heatmaps were constructed with Seaborn 0.12.1 in Python 3.10 and annotated with Inkscape (Rossum and Drake, 2009; Waskom and Seaborn development team, 2020). Toxin-antitoxin systems were identified with antiSMASH (Blin *et al*., 2021). CrisprCAS finder (Couvin *et al*., 2018) was used to establish the presence of cas systems in all extracted genomes.

## Results

### Genomes and Phylogeny

Four genomes were assembled from SRA data and an additional 22 recovered from the NCBI GenBank non-redundant database (Table 1). Of these 22 genomes, 16 came from environmental metagenome sequencing efforts that did not specifically target *Chlamydiota*, and the six that were targeted came from a single paper. Two were Identified to species and three to family level (Supplementary data 1). One of these, JAGXTH01, is a *Rhabdochlamydiaceae* misidentified as a *Simkania* sp. (see Figure 1 and accession GCA_018333475.1).

**Table 1.**
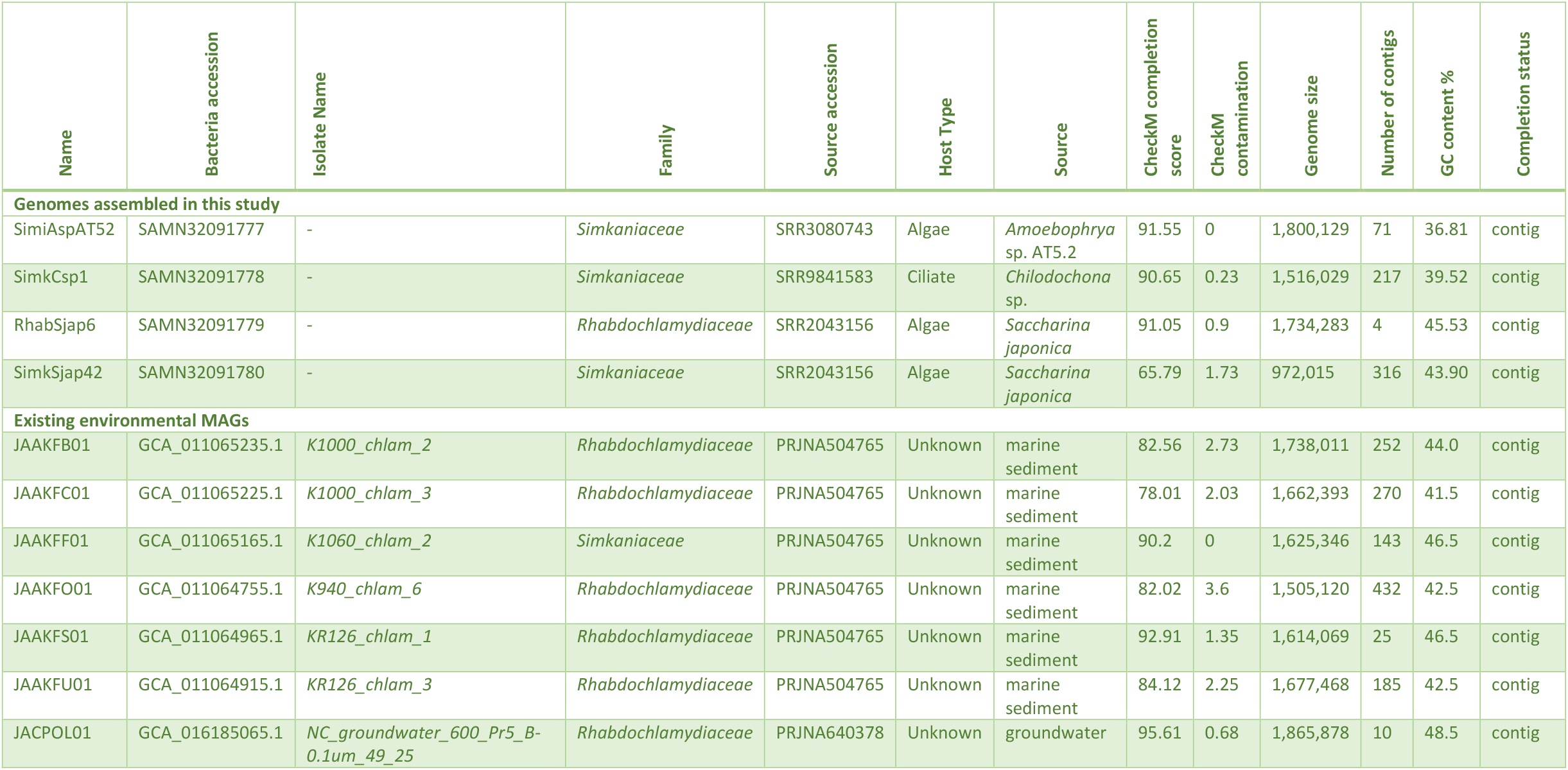
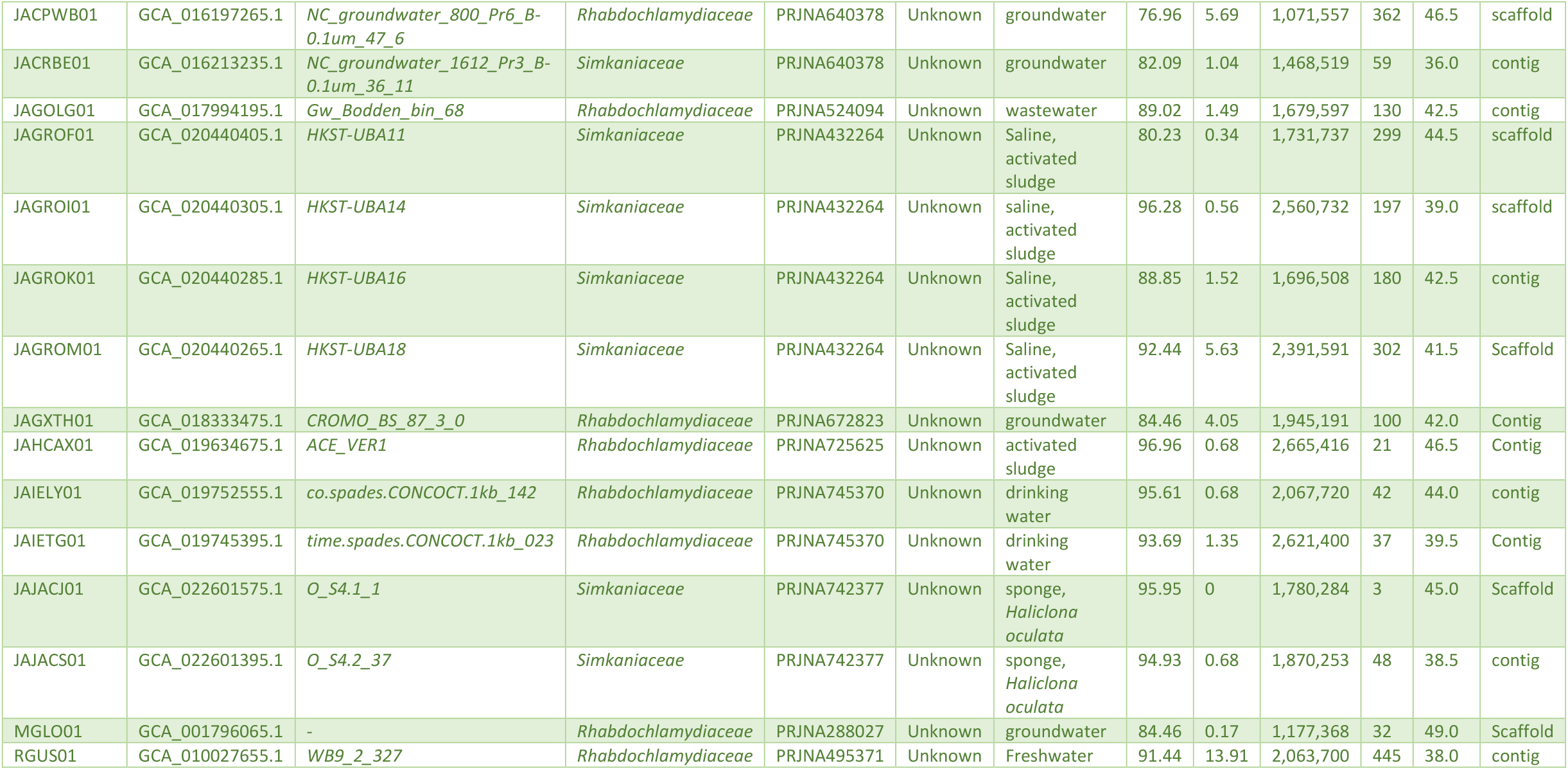
Select genome metadata for newly assembled Rhabdochlamydiaceae and Simkaniaceae genomes and previously assembled NCBI environmental MAGs. Full metadata can be found in Supplementary data 1.

**Figure 1.**
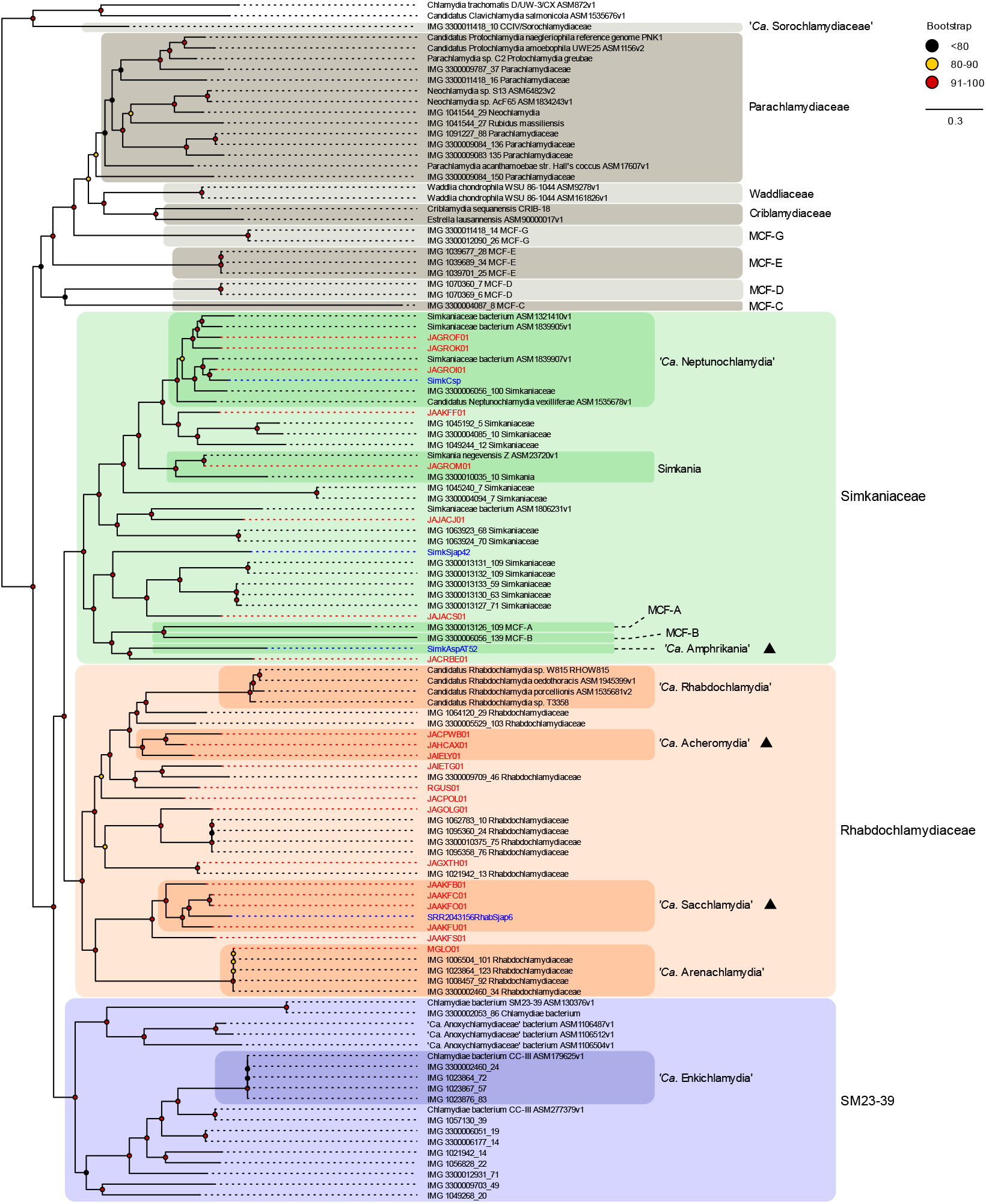
Genome wide phylogeny of Parachlamydiales. Maximum likelihood (ML) phylogeny of Parachlamydiales constructed from 34 single copy gene clusters that contain a total of 3604 genes. New genomes are indicated by ▲ with blue labels for genomes derived from host SRA bins, and red for MAGS without host assocition data. Bootstrap values based on 1000 replicates are indicated with coloured circles (red = 91-100, yellow = 81-90, black <= 80).

Several groups, like SM23-39 and most MCF types, are sufficiently different and diverse to warrant their own family name. In literature, SM23-39 has been referred to as both ‘*Ca*. Limichlamydiaceae’ and ‘*Ca*. Anoxychlamydiaceae’ (Pillonel, Bertelli and Greub, 2018; Dharamshi *et al*., 2020, 2023). Their classification seems to be mixed, with genomes identified as SM23-39 also spanning a second proposed family ‘*Ca*. Enkichlamydiaceae’ (Supplementary data 1 and Köstlbacher *et al*., (2021). Based on GTDBtk analysis we have opted for using the clade name SM23-39 to refer to this clade to avoid naming confusion, and because all genomes share family level classification by this method, as well as equivalent diversity with *Rhabdochlamydiaceae* and *Simkaniaceae* (Figure 1 and 2). There is no set threshold for classifying family or order level, and although GTDBtk does overclassify higher taxons (Chaumeil *et al*., 2020), it is currently the only objective way of designating family level classification. Higher classifications can be added later as needed, but at this stage we do not believe it is helpful to establish additional families or orders while the current within family phylogeny is so fluid.

**Figure 2.**
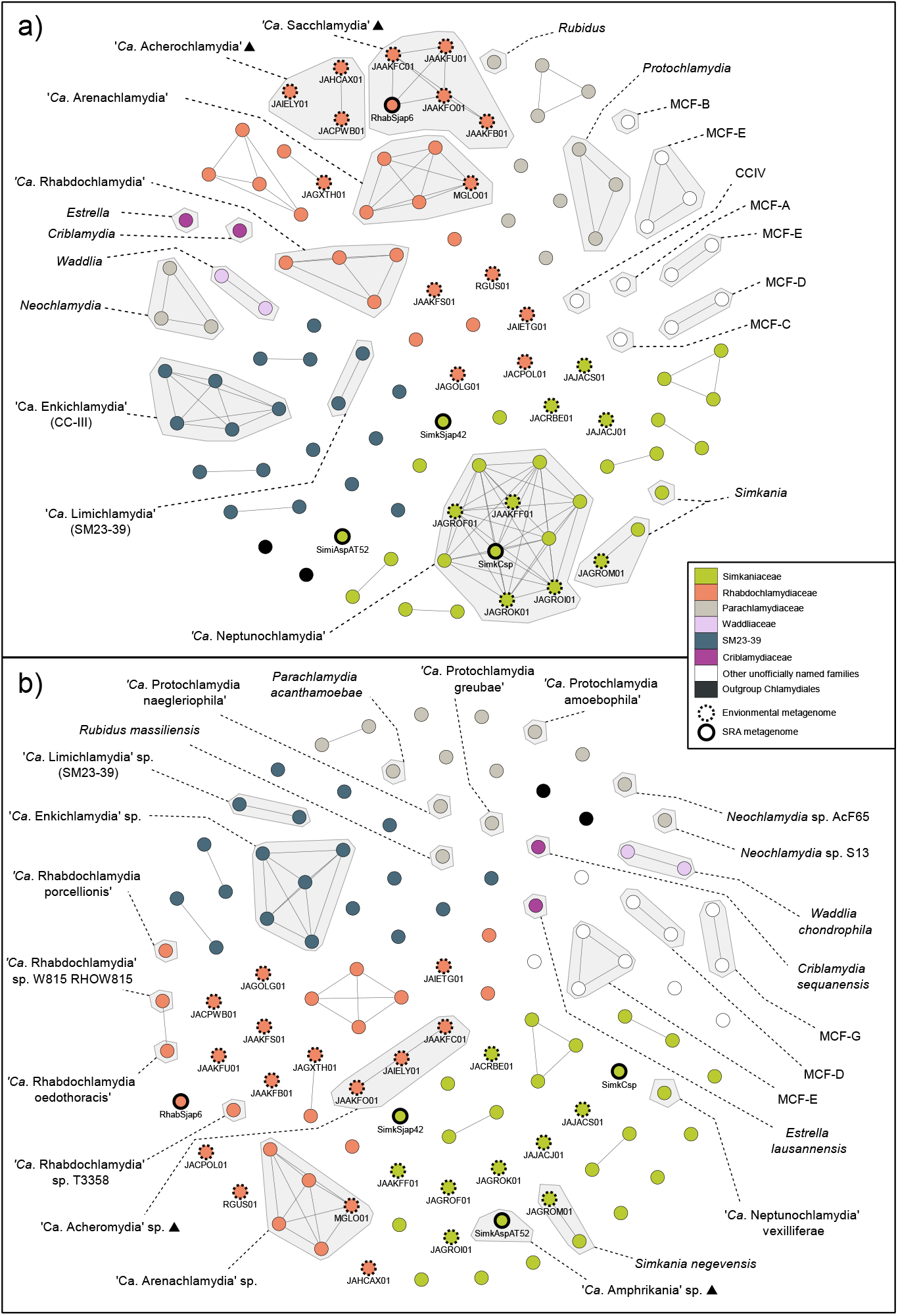
Genus and species level clustering across *Parachlamydiales*. Frutcherman Reingold networks of pairwise a) Average Amino Acid Identity (AAI) with edge weights >65% similarity and b) Average Nucleotide Identity (ANI) with edge weights >95% similarity across all genomes. AAI and ANI illustrate genus and species boundaries, respectively. Proposed genus names are indicated with a ▲

AAI and ANI results shown in Figure 2 support most existing genera and species clusters, but also indicates the potential need for some genera to be split (e.g., *Simkania*) and some species to be taxonomically combined (‘*Ca*. Rhabdochlamydia oedothora’ and ‘*Ca*. Rhabdochlamydia’ sp. W815 RHOW815).

Using AAI values with a cut off of 65%, we find objective support for naming the clade containing MGLO01 as its own genus, ‘*Ca*. Arenachlamydia’, as suggested in (Pillonel, Bertelli and Greub, 2018). However, contrary to Pillonel et al. (2018), we do not find that they are unique within *Chlamydiota* in lacking a Menaquinone biosynthesis/futalosine pathway (Supplementary data 1). Amongst the genomes included in this study, all lack the futalosine pathway (KEGG module M00930 and M00931) except for MCF-G and ‘*Ca*. Sorochlamydiaceae’ (clade CC-IV). The Menaquinone biosynthesis pathway (M00116) is partially present in all except 13 genomes, of which span several different clades across *Parachlamydiales* (Supplementary data 1). Other clades form putative genera groups of four or more genomes that are simply unnamed. *Simkaniaceae* and *Rhabdochlamydiaceae* in particular lack solid categorisation, each having only two previously named genera (Figure 1 and 2). Further, the genomic diversity displayed within both families is extensive. Some single genomes possess more than ten times more unique genes than genes shared across all genomes (Figure 3), though this could be an artifact of divergent orthologs not clustering together. It is very likely that *Parachlamydiales* will need to be split into multiple orders as more data becomes available, and revision of family level affiliation, to better describe this ancient clade of bacteria.

**Figure 3a.**
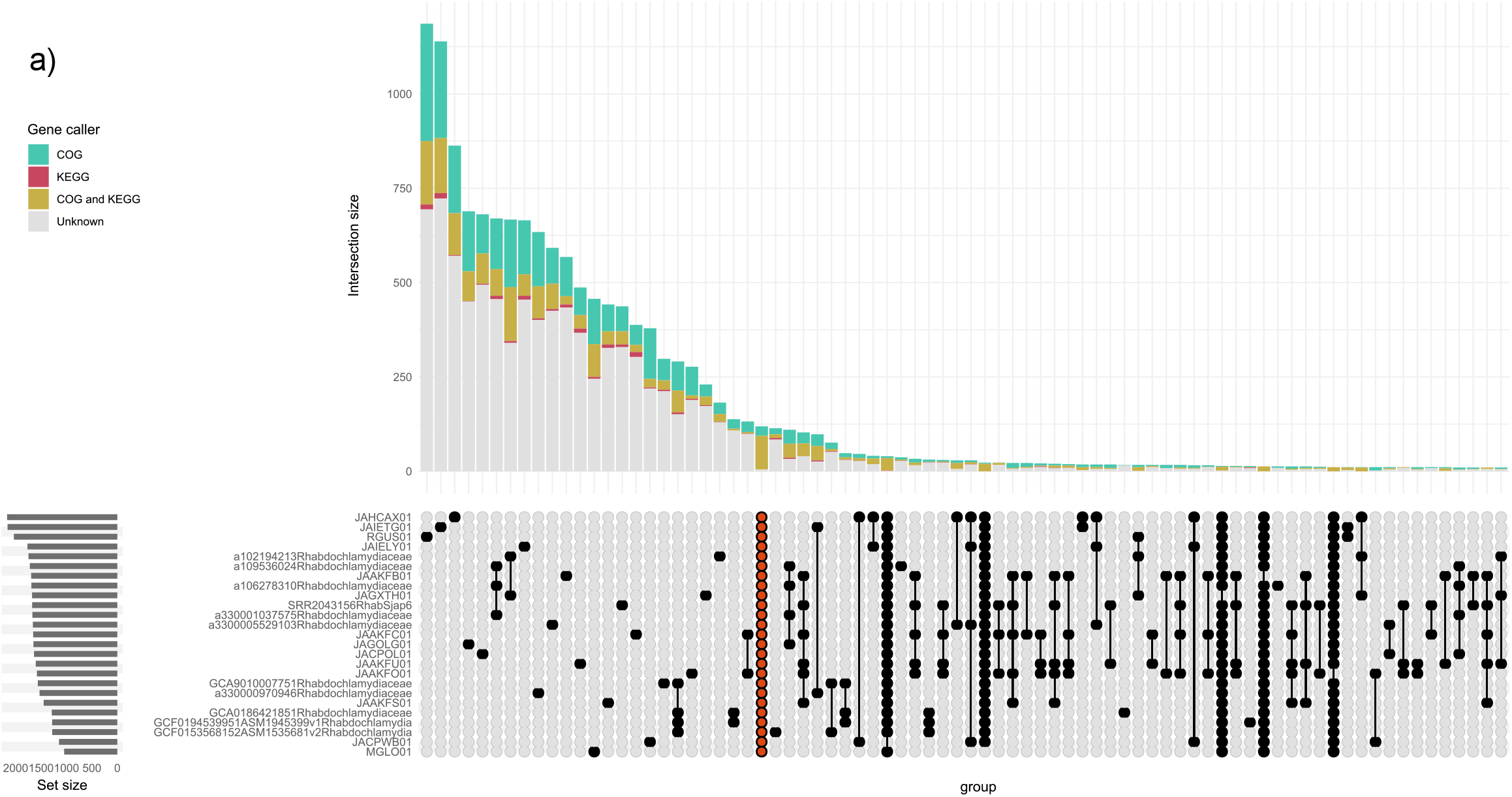
Gene content comparison for *Rhabdochlamydiaceae* and *Simkaniaceae*. An upsetplot illustrating the disparity of gene presence absence across the families a) *Rhabdochlamydiaceae* and b) *Simkaniaceae*. Vertical bars illustrate the number of genes present shared across compared genomes, indicated by black circles beneath each bar. Orange circles indicate the core genome shared by all genomes in the comparison.

**Figure 3b.**
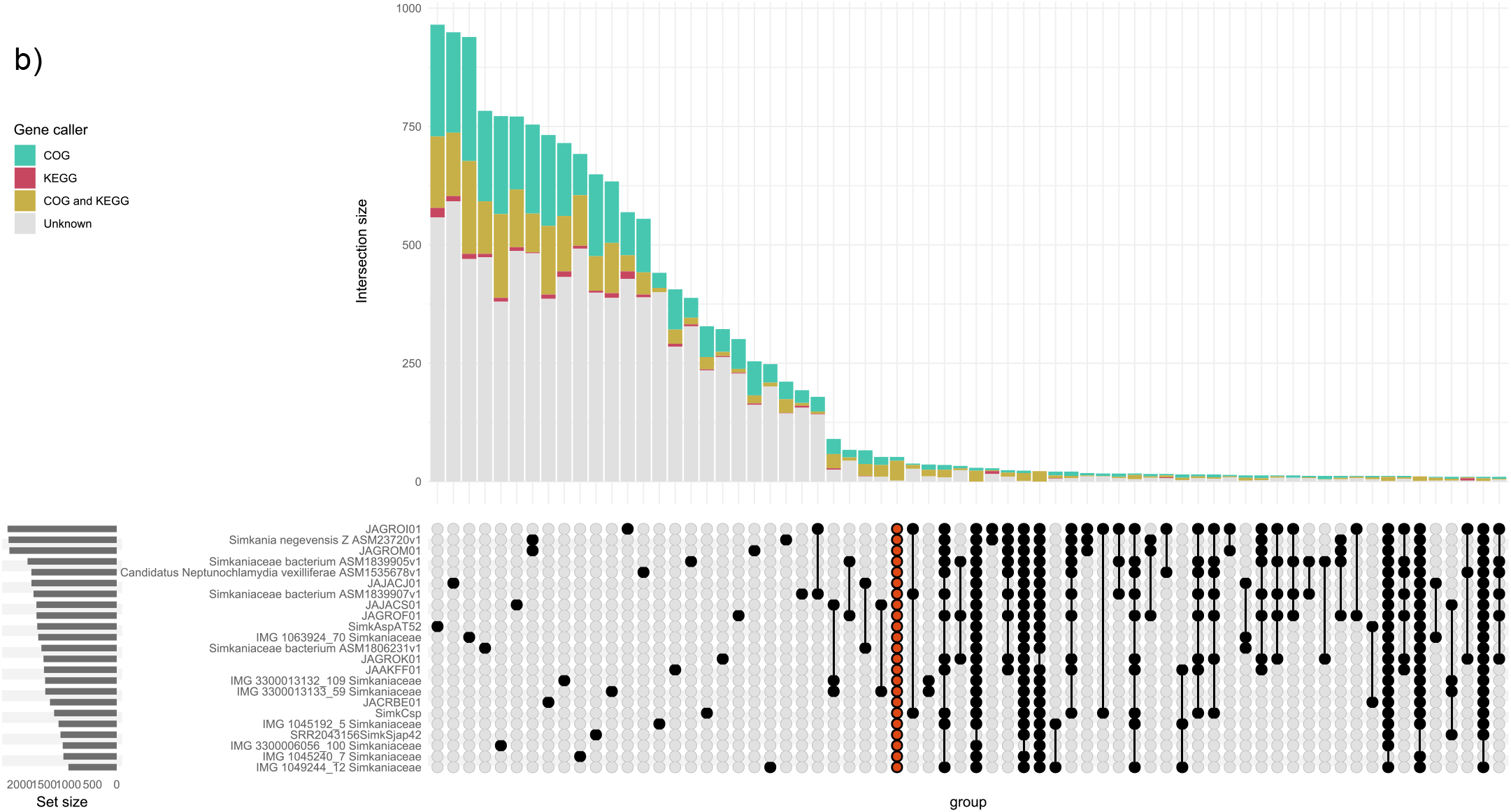
Gene content comparison for *Rhabdochlamydiaceae* and *Simkaniaceae*. An upset plot illustrating the disparity of gene presence absence across the families a) *Rhabdochlamydiaceae* and b) *Simkaniaceae*. Vertical bars illustrate the number of genes present shared across compared genomes, indicated by black circles beneath each bar. Orange circles indicate the core genome shared by all genomes in the comparison.

### Proposed Taxonomy

Based on AAI, ANI, GTDBtk analysis, metabolic results, and phylogeny, we find evidence for at least three new genus-level clusters within Rhabdochlamydiaceae and Simkaniaceae. The genera we propose contain either multiple high-quality environmental genomes found with the methods described in this study, or are high-quality assemblies extracted from potential host SRAs.

‘*Candidatus* Sacchlamydia’ gen. nov. - For the cluster that contains RhabSjap6, a host associated genome, named after the putative host *Saccharina japonica* and the pattern of using “-chlamydia” as the suffix for Chlamydiota genera.

‘*Candidatus* Amphrikania’ gen. nov. - A second new clade with host association data lies within the Simkaniaceae (the genome named SimkAspAT52). Amphri-after Amphritrite Queen of the Sea and echoing its marine host *Amoebophyra*, and -kania after the type genus for the family Simkaniaceae, *Simkania*.

Similarly to SimkAspAT52, clade SimkSjap42 is likely a new species group but will not be named because it has genome that is only 65% complete in CheckM analysis (Figure 1., Table 1).

‘*Candidatus* Acheromydia’ gen. nov. - For the clade including the MAGs JAHCAX01 and JACPWB01 which share >65% AAI similarity. For convenience, and so as not to over inflate this group with new names, I also include the relatively closely related genome JAIEL01 (>62% AAI similarity). This genus is named after Acheron (Acher-) one of five Greek rivers of the underworld that occasionally surfaces above ground. The MAGs come from drinking water, activated sludge and ground water. -mydia from *Chlamydia*.

### Metabolism

Parachlamydiales have a broad array of metabolic pathways (Figure 4 and Supplementary data 1), including nutritional and defensive systems. Of note, several have mostly complete B vitamin synthesis pathways (biotin, riboflavin and thiamine), known to be associated with beneficial contributions to the host in other symbioses with intracellular bacteria.

**Figure 4.**
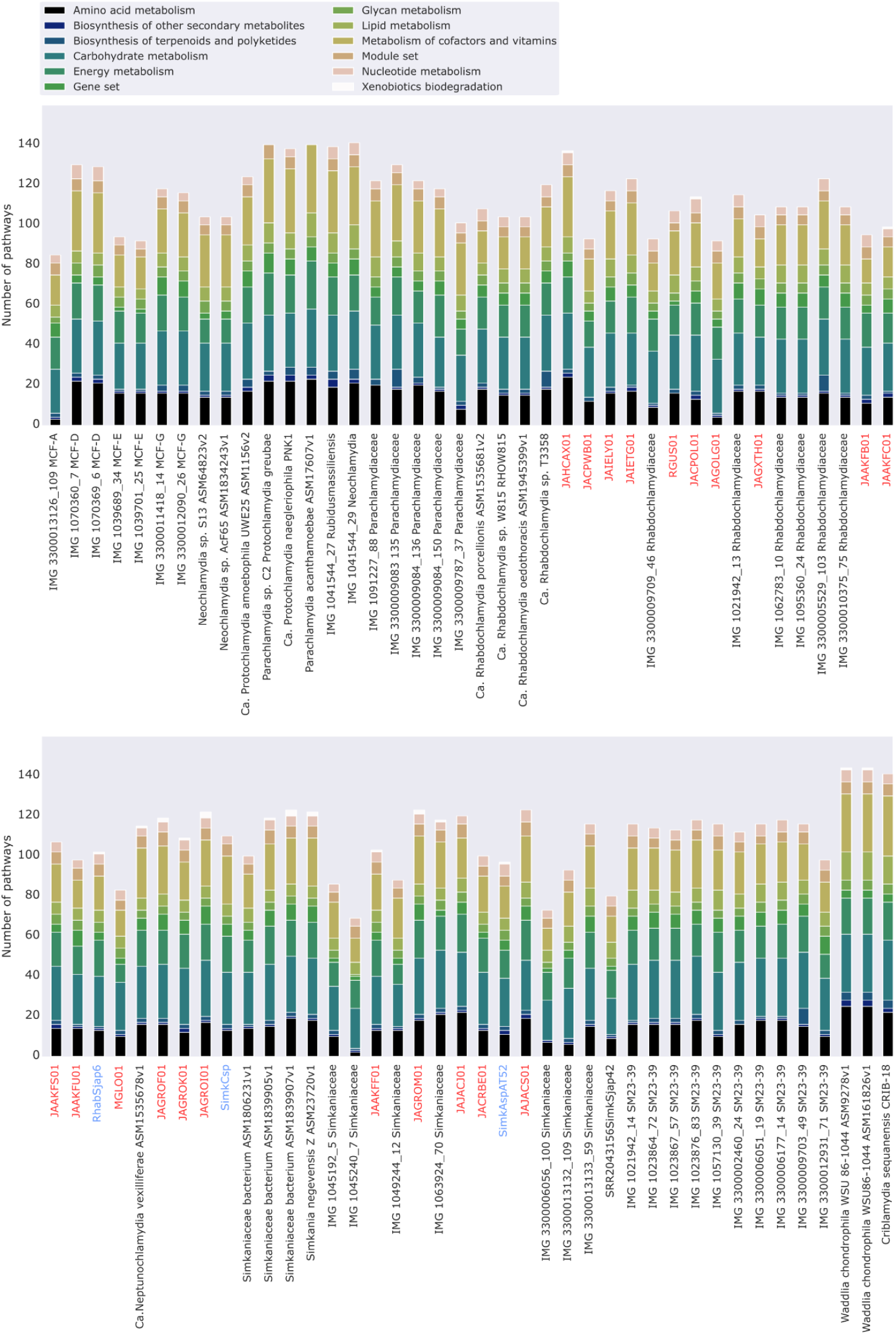
The number of each type of metabolic pathway found in each *Chlamydiota* genome. Red indicates environmental MAGs; blue indicates MAGs assembled in this study.

Some Parachlamydiales have uroporphyrinogen decarboxylase and uroporphyrinogen-III synthase which form part of several biosynthesis pathways including heme, chlorophyll, cobalamin, and siroheme. In addition, some have incomplete phylloquinone (vitamin K1) biosynthesis pathways. All Parachlamydiales genomes here also have complete Type III secretion systems and incomplete gene sets associated with flagellar assembly (Supplementary data 1). Several terpenoid synthesis pathways are also present, including dTDP-L-rhamnose biosynthesis.

We find that KEGG Kofam identifies CRISPR-associated protein Cas2 in all *Parachlamydiales* studied in this study, including *P. acanthamoeba*. Several have additional types of CRISPR/Cas associated genes (Supplementary data 1). CRISPRcas-finder identifies cas and cas spacers in the four MAGs extracted from the SRA samples (Supplementary data 1) and only RhabSjap6 appears to possess both CRISPR elements and a cas cluster. Whether these genes are functional or associated with defensive symbiosis against phages based purely on metabolic prediction is unknown; the class of cas is not predictive of their function.

## Discussion

### Genomes and Phylogeny

We observe high genomic, phylogenetic, and metabolic diversity in the currently named families, genera, and within species of *Chlamydiota*, in line with other studies (Horn, 2008; Lagkouvardos *et al*., 2014; König *et al*., 2017). The current taxonomy is not sufficient and has been left undefined to the detriment of these bacteria, in part due to the nature of the clade being so broad and data being so sparse. This has resulted in some genomes such as *Rubidus massiliensis* and JAGXTH01 being miscategorised on NCBI. *Rubidus massiliensis* is categorised as *Chlamydiaceae* rather than *Parachlamydiaceae*, as proposed in the study where it was first described (Bou Khalil *et al*., 2016) and as shown in Figure 1. MAG JAGXTH01 is identified as a *Simkania* sp., whereas we find it falls within the *Rhabdochlamydiaceae*.

Prior to this study, only two Environmental MAGs found had been assigned a taxonomic classification lower than family, and one of those two is misidentified (Supplementary data 1). At this stage we do not believe it is helpful to split existing family clades like SM23-39 into multiple families, especially since they are supported by GTDBtk analysis. There is also some confusion in literature for the taxonomy for some clades: ‘*Ca*. Limichlamydiaceae’ (Pillonel, Bertelli and Greub, 2018) is included in ‘*Ca*. Anoxychlamydiaceae’ (Dharamshi *et al*., 2023), and ‘*Ca*. Enkichlamydiaceae’ (Pillonel, Bertelli and Greub, 2018) == Chlamydiae clade III (Dharamshi *et al*., 2023).

We believe the splitting of *Parachlamydiales* into three new orders are in some respects warranted: ‘C*a*. Anoxychlamydiales’, ‘C*a*. Simkaniales’ (which includes *Rhabdochlamydiaceae* and *Simkaniaceae*) and ‘C*a*. Amoebachlamydiales’ (for *Parachlamydiaceae*) (Dharamshi *et al*., 2023). We could also argue that *Rhabdochlamydiaceae* and *Simkaniaceae* will eventually need their own Order classifications based on the high gene content diversity within the families (figures 3a and 3b). However, we do not believe it is helpful to split *Parachlamydiales* into further orders at this time while most families and genera have not been properly defined. Additionally, Order classification has no robust criteria and the erection of further Orders may contribute to confusion around the taxonomy of *Chlamydiota*.

### Metabolism

Vitamin B biosynthesis is known to be associated with beneficial contributions to the host in other symbioses with intracellular bacteria. However, all characterised Parachlamydiales have biphasic lifestyles, and many are known pathogens, so the presence of B vitamin modules does not necessarily indicate potential benefit to the host. For instance, pathogenic *Chlamydia* with bioY genes have previously been observed to uptake biotin from host cells (Fisher *et al*., 2012).

If Parachlamydiales do form beneficial symbioses with their host it is feasibly associated with photosynthesis through vitamin Phylloquinone synthesis or chlorophyll synthesis. Phylloquinone is produced and used by plants, algae, and cyanobacteria in photosynthesis. Most *Parachlamydiales* genomes also have a complete chorismate synthase pathway (Figure 5, Supplementary data 1), an important intermediate product for phylloquinone production, as well as intermediates for alkaloids and salicylic acid that are important in plant defence systems (Hamberger *et al*., 2006; Shanmugabalaji *et al*., 2022). It should be noted that chorismate can also be used to synthesize a variety of other vital aromatic compounds such as ubiquinone, which is common in bacteria (Dosselaere and Vanderleyden, 2001).

**Figure 5.**
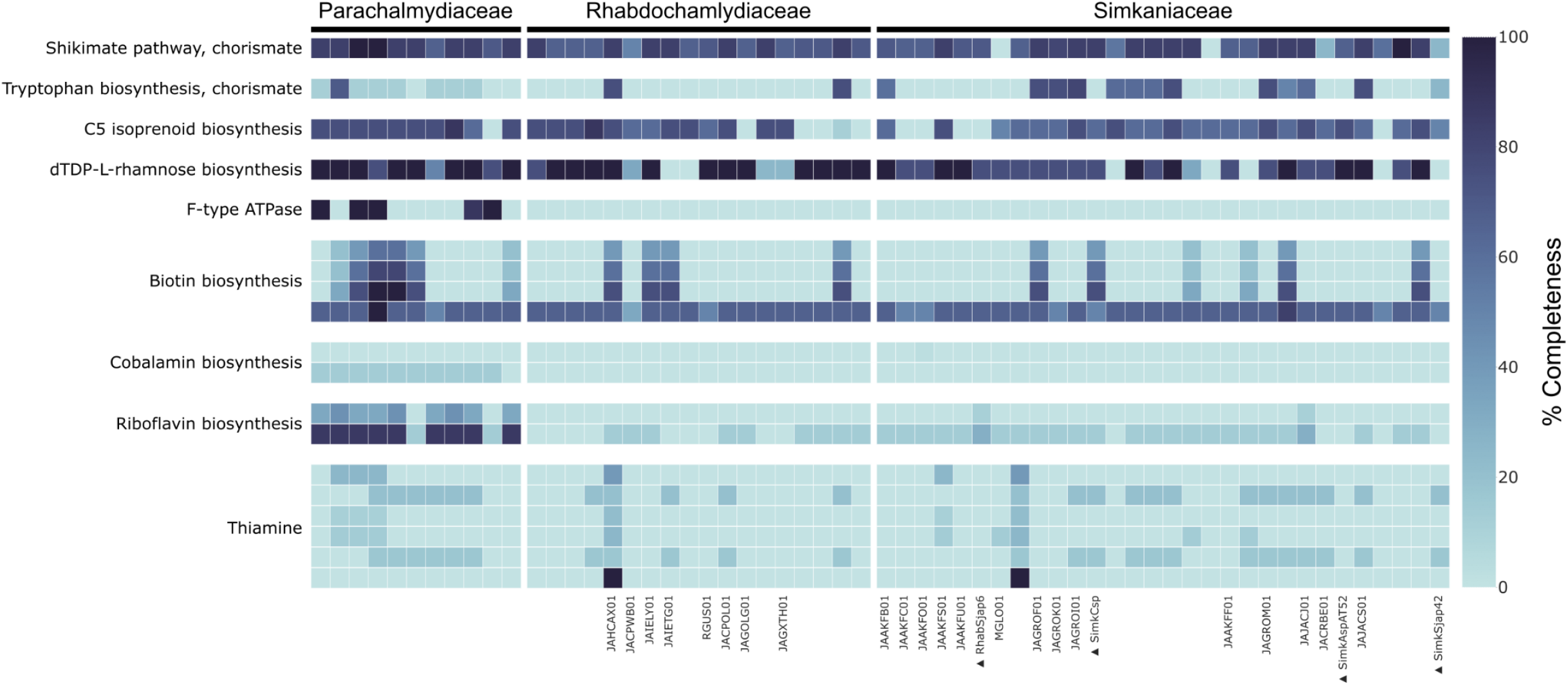
Heatmap of the completeness of KEGG metabolic pathways of interest across the *Parachlamydiaceae, Rhabdochlamydiaceae* and *Simkaniaceae*. Full metabolic pathway completeness for all genomes and all other pathways are available in Supplementary data 1. New genomes assembled in this study are indicated by ▲

There is also a possibility that these bacteria form defensive symbioses, as seen in the case of viral protection of amoebae by *P. acanthamoeba* (Arthofer *et al*., 2022). Unfortunately, we do not know how *P. acanthamoeba* blocks viral factory formation, so we cannot identify any associated pathways through homology. The presence of CRISPR/cas systems in some genomes could indicate a potential mechanism for this trait, but we do not know that they are functional, and, if they are, they may be associated with the free living stage rather than antiviral activity in the host. CRISPR/Cas Phage defence are common across bacteria but notably rare in microbial symbionts (Deveau, Garneau and Moineau, 2010). They have also previously been found in *Protochlamydia* genomes and other *Chlamydiota* (Deveau, Garneau and Moineau, 2010; Bertelli *et al*., 2016; Köstlbacher *et al*., 2021).

An alternative route for defensive symbiosis could be terpenoid production. Terpenoids such as dTDP-L-rhamnose are associated with plant-bacteria symbioses as well as pathogenicity (Ma, Pan and McNeil, 2002; Jofré, Lagares and Mori, 2004; French, 2017; Jiang *et al*., 2021). Terpenoids are also used by algae themselves in defence against threats such as bacteria and heavy metals (French, 2017; Karimi *et al*., 2019). Some Red algae use bacteria-like pathways to produce their terpenoids (Wei *et al*., 2019).

## Conclusions

Chlamydiota are hyper diverse and under described. We have added to the known phylogenies of the *Parachlamydiales* families: *Simkaniaceae* and *Rhabdochlamydiaceae*. In addition, we clarify the status of two additional *Rhabdochlamydiaceae* and one *Simkaniaceae* genera, for which we propose the names ‘*Ca*. Sacchlamydia’, ‘*Ca*. Acherochlamydia’, and ‘*Ca*. Amphrikania’. The metabolic potential of *Chlamydiota* is likewise highly diverse, with several clusters of genes that could be associated with both defensive and nutritional symbiosis or otherwise associated with pathogenicity. Closer study of the metabolic interactions between *Parachlamydiales* and their microeukaryotic hosts are required to elucidate if and how they affect each other.

## Supporting information

Supplementary data 1 - Metadata

Supplementary data 2 - Partition model

## Abbreviations

MAG: Metagenome Assembled Genome
ANI: Average Nucleotide Identity
AAI: Average Amino-acid Identity

## Acknowledgements

Conceptualisation, supervision, methodology and funding acquisition – HRD and GDDH

Investigation –HRD

Analysis, Visualisation, and Data Curation – HRD

Original Draft Preparation – HRD and GDDH

Additional thanks to Stefanos Siozios for guidance early on in this project.

## Supplementary material captions

Supplementary data 1 – metadata and metabolic data for genomes used in this study.

Supplementary data 2 – tree partition models for Figure 1.

